# Modeling environmental surveillance of *Dracunculus medinensis* in aquatic habitats using a three-dimensional agent-based model

**DOI:** 10.64898/2026.05.05.722897

**Authors:** Jaewoon Jeong, Rebecca Garabed

## Abstract

Guinea worm disease eradication efforts may benefit from environmental surveillance methods capable of detecting infected copepod intermediate hosts in aquatic habitats. We developed a three-dimensional, spatially explicit agent-based model to examine how ecological processes influence detection probability for a hypothetical water sampling method. The results show that surveillance sensitivity is shaped by the combined effects of larval diffusion, copepod density, and pond size, with interactions among these factors producing nonlinear relationships. Detection, in our model, was concentrated within a relatively restricted period after larvae matured to the infective stage and before dispersal and mortality reduced presence, indicating a limited spatiotemporal window for effective sampling. Surveillance performance peaked under intermediate dispersal regimes that generated sufficient spatial overlap between larvae and intermediate hosts, while both limited dispersal and excessive diffusion reduced detection by constraining encounters or diluting larval concentrations. Increasing habitat size reduced detection by diluting larval concentrations, but the magnitude of this effect depended on copepod density and dispersal dynamics, producing nonlinear and threshold responses rather than simple scaling with pond volume. Spatial and temporal patterns of detection shifted as larvae dispersed, with the most favorable detection periods occurring when both larval abundance and intermediate host encounters were elevated. These findings indicate that surveillance can be guided by local ecological conditions. When the timing of larval introduction is uncertain, effective surveillance requires repeated sampling over time to capture transient windows of detectability and the sampling will be less effective in very stagnant and highly mixed waterbodies. Overall, this study demonstrates how mechanistic modeling can support the design and interpretation of environmental surveillance strategies for Guinea worm eradication programs.

**Author summary:** Guinea worm disease is close to eradication but confirming that transmission has fully stopped remains difficult because detecting infectious larvae in water is challenging. Transmission depends on freshwater copepods that become infected after ingesting Guinea worm larvae. These copepods are short-lived and unevenly distributed within ponds, and infected individuals may die before larvae reach the infective stage. As a result, environmental detection is inherently uncertain. We developed a three-dimensional agent-based model to simulate larval dispersal, copepod infection, and water sampling in a pond environment. The model shows that detection is constrained to a brief period when mature larvae and copepods overlap in space and time, and that this window depends strongly on local ecological conditions such as larval dispersal, copepod density, and pond size. Because infected copepods can be present outside these narrow detection windows, negative water samples do not necessarily indicate absence of transmission, highlighting the need for repeated, spatially targeted surveillance during the final stages of eradication.

## Introduction

Guinea worm disease (dracunculiasis), caused by the parasitic nematode *Dracunculus medinensis*, is a water-based disease that once affected millions of people across Africa and Asia (1). Transmission occurs when individuals consume untreated water containing copepods infected with third stage larvae (L3s). Inside the definitive host, the larvae mature and mate, and after about a year, gravid female worms migrate to the skin and emerge through painful ulcers typically on the foot, often leading to secondary infections and long-term disability (2). There is currently no vaccine or curative drug for the disease, and prevention relies entirely on safe water access, containment, behavior change, and surveillance. While diagnostic assays for the parasite have been developed and continue to advance, none are yet routinely deployed or operationalized for environmental surveillance. Coordinated efforts led by The Carter Center, WHO, other partners of the Guinea Worm Eradication Program, and Ministries of Health in Guinea worm endemic countries have resulted in dramatic decline in Guinea worm cases from millions in the 1980s to a few dozen annually in recent years (3). However, eradication has been hindered by the emergence of additional definitive hosts—particularly dogs—and the persistent difficulty of detecting residual transmission in low-endemic settings (4). As the global eradication program approaches the endgame phase, sensitive and operationally feasible surveillance tools are increasingly important to identify residual transmission and prevent re-establishment of the parasite.

Copepods, a type of freshwater zooplankton, serve as essential intermediate hosts in the life cycle of *D. medinensis*, ingesting first-stage larvae (L1s) that then develop into infective L3s. Among cyclopoid copepods, genera such as *Mesocyclops* and *Thermocyclops* have been identified as competent intermediate hosts for *D. medinensis* due to their habitat preferences and ingestion behavior (5,6). However, several biological features of the host-parasite interaction limit transmission efficiency. Only adult copepods are capable of harboring L3s, and infected individuals often experience high mortality before larval development is complete (5,7).

Moreover, successful transmission to definitive hosts requires ingestion of multiple infected copepods, since single ingestions rarely result in establishment of both male and female worms (4). These ecological and transmission constraints likely reduce the density of infective copepods in natural water bodies, making environmental detection particularly difficult in low-transmission settings. Consequently, understanding the spatial and temporal dynamics of infected copepods may help improve surveillance strategies aimed at identifying contaminated water sources and interrupting residual transmission.

Environmental surveillance for Guinea worm through detection of copepods infected with infective L3 larvae could be transformational in identifying high-risk water sources, but several challenges exist. Field data show that the prevalence of infected copepods in endemic areas is typically low, with reported averages around 5.2% with a range from 0.5% to 33.3% (4).

Furthermore, infected copepods may not survive the full developmental period required for larvae to reach the infective L3 stage, as infection often leads to premature mortality in addition to background natural mortality (5), thereby reducing the probability of detecting infective larvae in environmental samples. In terms of spatial ecology, *Mesocyclops*—a common intermediate host—exhibit non-random distributions, with a tendency to aggregate near pond margins and migrate vertically in response to environmental cues such as oxygen levels and predation pressure (8,9). These behaviors suggest that infected copepods may remain spatially clustered rather than evenly dispersed, further limiting the likelihood of their detection during standard water sampling efforts. Together, low infection prevalence, infection-associated mortality, and spatial aggregation of copepods create substantial challenges for environmental surveillance in Guinea worm endemic areas.

Detecting copepods infected with *Dracunculus medinensis* in aquatic environments poses a significant challenge for Guinea worm surveillance. Field detection is limited by low infection prevalence, brief infectivity periods, and spatial heterogeneity in host and parasite distributions (4,5). These constraints reduce the effectiveness of empirical sampling alone. Mechanistic modeling can complement field surveillance by simulating how ecological and spatial processes influence the probability of detecting infected copepods under different environmental conditions. For example, a previous study (10) used a spatially structured model of schistosomiasis to examine how environmental variation, snail host ecology, and hydrological patterns influence long-term transmission outcomes. This highlights how aquatic models can guide surveillance design and improve targeting of control efforts. Agent-based models (ABMs) are particularly well-suited for this task, as they represent individual copepods with spatial positions, life histories, and infection status. When integrated into three-dimensional (3D) aquatic environments, ABMs can incorporate key processes such as larval diffusion, copepod movement, and the spatial configuration of water sampling strategies (11). By linking ecological processes with surveillance performance, these models can help identify conditions under which environmental detection is most likely to succeed and inform the design of operationally feasible sampling strategies for Guinea worm eradication programs.

The objective of this study was to develop a 3D, spatially explicit, ABM to simulate the dynamics of *D. medinensis* larvae and their interactions with copepod intermediate hosts in a pond ecosystem. The model incorporated key processes—larval diffusion, ingestion and infection within copepods, infection-induced mortality, and water sampling for surveillance. Although specific environmental drivers of larval diffusion, such as water temperature, turbulence, and vegetation structure, were not modeled explicitly, variation in the larval diffusion rate was used as parsimonious representation of their combined influence on larval dispersal. By systematically varying this larval diffusion parameter, copepod densities, and pond size, we estimated the probability of detecting infected copepods under a range of ecologically plausible conditions. We first established model behavior using baseline parameter values, then performed sensitivity analyses to identify which factors most strongly influenced detection probability and to elucidate potential interactions among parameters. These analyses were used to evaluate how ecological variability may influence surveillance performance across different aquatic environments. Overall, this mechanistic modeling framework provides insight into ecological and sampling conditions that may improve environmental detection of Guinea worm transmission and supports the development of operationally relevant surveillance strategies for ongoing eradication efforts.

## Methods

### Study system and model overview

We developed a 3D, spatially-explicit, ABM to simulate the ecological and transmission dynamics of *D. medinensis* larvae and their interactions with cyclopoid copepod intermediate hosts in a closed pond ecosystem. The model captured key processes relevant to environmental surveillance, including larval dispersal, copepod movement and behavior, infection progression, and detection probability. Each copepod was represented as an individual agent with unique spatial coordinates, age, and lifespan, while larval densities were tracked across a three-dimensional grid. Copepod movement followed a correlated random walk modulated by edge-biased spatial preferences, and larval diffusion was modeled as a stochastic process determined by a specified diffusion coefficient. The simulation advanced in daily timesteps, incorporating the dispersal of L1s, random movement and aggregation of copepods, larval encounter and ingestion events, and within-copepod larval development. Ingested L1s developed to the infective L3 stage within copepods, with infection-induced mortality increasing if larval burden exceeded a defined threshold. Background mortality and recruitment of new adult copepods were included to maintain population turnover within the simulated pond environment (Fig 1). Water sampling was implemented as a spatially and temporally explicit process, enabling estimation of the probability of detecting L3-infected copepods under a range of ecologically plausible surveillance scenarios.

**Fig 1.**
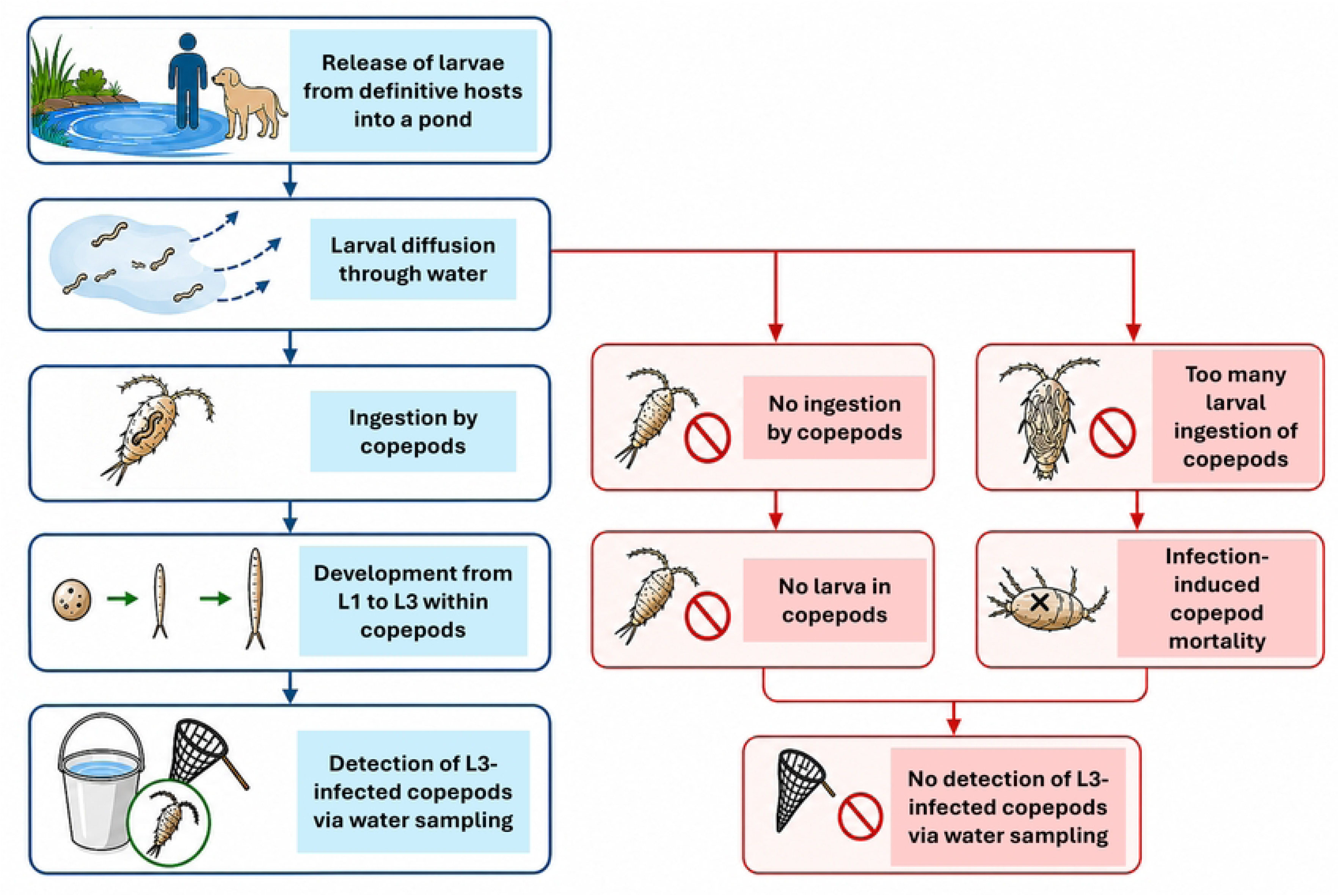
Conceptual framework of Guinea worm transmission dynamics and environmental surveillance outcomes. The diagram illustrates alternative ecological pathways following release of *Dracunculus medinensis* larvae into a pond, resulting in either successful detection or non-detection of L3-infected copepods through environmental water sampling.

### Aquatic environment

The aquatic environment was modeled as a hypothetical cylindrical pond (Fig 2), with the three-dimensional spatial domain discretized into cubic grid cells at a 10-centimeter resolution, each representing a 1-liter volume. The x–y–z Cartesian coordinate system was used to track the positions of individual copepods and larvae, allowing for detailed simulation of their spatial distribution and movement throughout the water column. This fine-scale structure enabled realistic modeling of larval diffusion, copepod aggregation, and spatial heterogeneity relevant to environmental sampling. The total pond volume, determined by the specified geometric parameters, was used to calculate the copepod population size for each simulated density condition.

**Fig 2.**
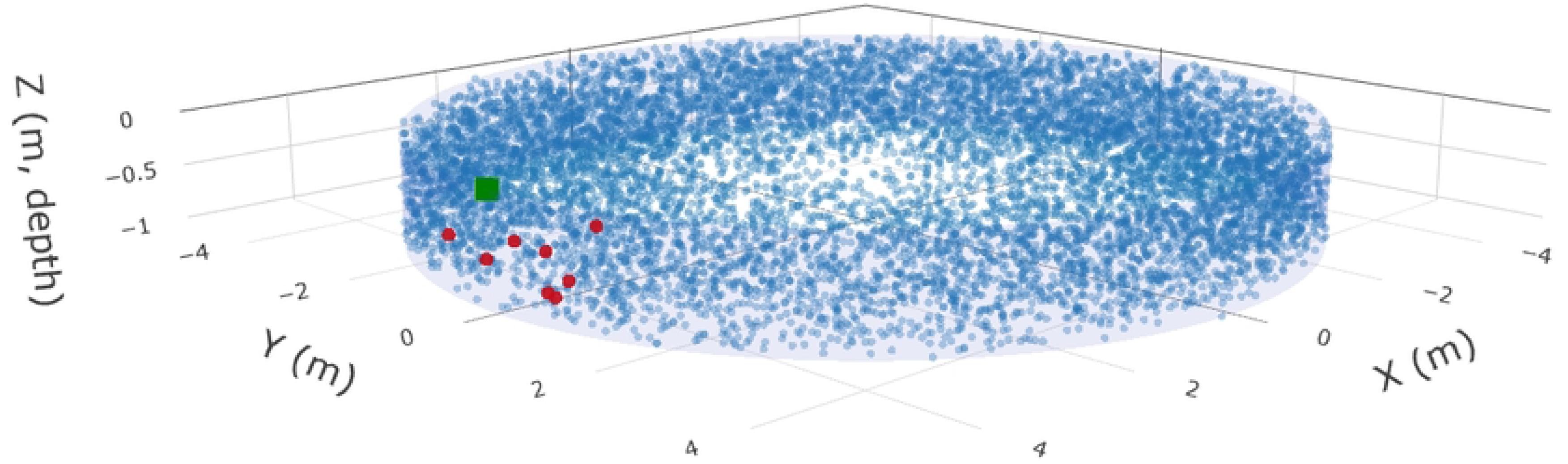
Three-dimensional spatial distribution of copepods in the simulated pond at a representative timestep on day 22. Red points indicate copepods carrying at least one L3-stage larva, blue points are uninfected copepods, and the green point marks the larval release site. L3-infected copepods are concentrated at intermediate distances from the release point, reflecting the combined effects of larval dispersal, copepod movement, and infection-induced mortality near the release area. Interactive three-dimensional image is available at https://rpubs.com/jwjeong/1411531.

### Larval Diffusion

At the start of each simulation, a fixed number of first-stage (L1) *D. medinensis* larvae (*N*_0_ = 2,000,000) was released at the pond’s surface edge *(R, 0, 0)* (1,2). As the simulation progressed, larval densities within each grid cell were updated at each timestep. Larval movement was modeled as a passive process, governed by three-dimensional Fickian diffusion originating from the point of release (11,12). The spatial distribution of larvae at location (*x*,*y*,*z*) and time *t* was defined by the analytical solution to the diffusion equation:

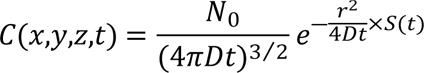 where *C*(*x*,*y*,*z*,*t*) is the larval concentration at position *(x, y, z)* and time *t*, *r* is the Euclidean distance from the release point, *D*_*c*_ is the diffusion coefficient (m^2^/day), and *S*(*t*) is the proportion of larvae surviving at time *t*.

The diffusion coefficient *D*_*c*_ was estimated as *D*_*c*_ = *v*^2^*τ*/*n* using empirical measurements of larval swimming, where *v* = 0.1567 *mm*/*s* is the average L1 swimming speed (13), *τ* = ^5^ is the reorientation time (s) (14), and *n* = 3 represents the three spatial dimensions. While this calculation reflected self-propelled movement in still water, natural aquatic environments may exhibit turbulence and microcurrents that increase larval dispersal beyond that predicted from swimming behavior alone.

Based on the average lifespan of 5 days *T*_*L*_ for free-floating L1 larvae, L1 larval survival time in the baseline model was treated as a random variable drawn from a PERT distribution (min = *T*_*L*_/ 2, mode = *T*_*L*_, max = *T*_*L*_× 2) (2).

### Copepod infection with Guinea worm larvae

We modeled *Mesocyclops* copepods as autonomous agents, explicitly tracking their spatial positions, infection status, and internal larval burden over time. The model included only adult copepods, as they are the primary life stage capable of ingesting and supporting the development of *D. medinensis* larvae to the infective L3 stage (2). Juvenile stages were not modeled explicitly; instead, a constant background recruitment process represented the steady maturation of juveniles into the adult population under equilibrium conditions. This assumption allowed the model to focus specifically on the subset of copepods most relevant to Guinea worm transmission and environmental detection.

Each copepod was assigned a lifespan drawn from a uniform distribution between 14 and 42 days (8,15) and randomly positioned within the three-dimensional spatial domain. Horizontal locations were generated by sampling an angle *θ*∼*U*(0, 2*π*) and a radial distance 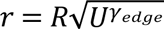, where *U*∼*U*(0, 1). *R* is the pond radius, and *γ*_*edge*_ controls spatial bias toward the pond edge.

This approach allowed simulation of edge preference behavior, with *γ*_*edge*_ = 1 yielding a uniform spatial distribution and *γ*_*edge*_ = 0 concentrating all copepods at the pond edge. Coordinates were calculated as *x* = *r*cos(*θ*), *y* = *r*sin(*θ*), and *z* was sampled uniformly between 0 (surface) and the maximum pond depth.

At each daily timestep, copepods encountered larvae based on the local larval density interpolated from a precomputed 3D grid. For copepod *j* at time *t*, the expected larval encounter rate *λ*_*j*_(*t*) was calculated as

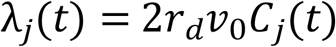

where *r*_*d*_ = 0.35 *mm* is the detection radius (16), *v*_0_ = 1*mm*/*s* is the swimming speed (17,18), and *C*_*j*_(*t*) is the larval concentration at the copepod’s current location. This formulation follows Gerritsen & Strickler (19) and assumes each copepod sweeps a cylindrical volume of water per unit time.

The ingestion rate for copepod ^j^ at time ^t^ was defined as the product of the larval encounter rate and the probability of ingestion upon encounter: *v*_*j*_(*t*) = *λ*_*j*_(*t*) × *P*_*ingest*_. To account for reduced ingestion as internal larval burden increases, the ingestion probability *f*_*ingest*_, was modeled using a logistic function: 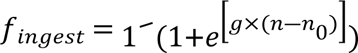, where *n* is the copepod’s current larval burden, *n*_0_ = 4 is the inflection point at which ingestion drops to 50%, and *g* = 2 controls the steepness of decline (5,7). This formulation represents a saturating behavioral response—copepods with few larvae ingested readily, but ingestion declined sharply once burden approached the threshold. The expected number of larvae ingested by copepod *j* at time *t* was therefore

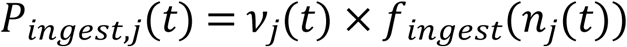

with the actual number drawn from a Poisson distribution with mean *P*_*ingest*,*j*_(*t*). Ingested larvae developed from L1 to L3 over a fixed period of 14 days (20).

Copepod mortality included two components. Age-dependent background mortality was applied at a constant daily rate of 1/30 (21), with individuals dying when their age exceeded their assigned lifespan. Infection-induced mortality was modeled such that survival probability declined with increasing larval burden according to:

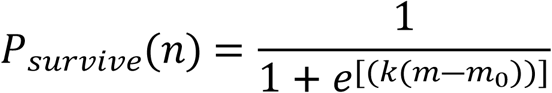

where *m* is the total number of ingested larvae at the timestep, *m*_0_ = 4 is the inflection point for 50% survival, and *k* = 1 controls the steepness of the decline (5,7). This logistic relationship captured the increasing mortality risk associated with high parasite burden (22).

When a copepod died of age-related causes, it was replaced by a newly recruited adult (age = 0) with a randomized lifespan and spatial position, maintaining a constant baseline population through recruitment from unmodeled juvenile stages. In contrast, copepods that died from infection were not replenished, allowing the adult population to decline in response to parasite-induced mortality. This approach ensured demographic stability under background mortality while capturing infection-driven reductions in copepod abundance.

### Water Sampling and Detection Probability

To evaluate surveillance sensitivity, we simulated water sampling to detect copepods infected with L3 *D. medinensis* larvae. At each designated sampling time point, a fixed number of water samples were collected by randomly selecting sphere-shaped volumes within the pond. Sample locations were generated using the same three-dimensional Cartesian coordinate system and spatial resolution as the model grid, ensuring consistency with the simulated distributions of copepods and larvae. Each water sample was defined as a sphere with a fixed volume of 24 liters, corresponding to standard field protocols used for Guinea worm surveillance. Sampling was implemented as an observational process at a frozen point in time that did not alter copepod behavior or water dynamics; disturbance effects associated with human entry into the water and removal of water with subsequent redistribution of water to fill the hole were not represented.

For each sampling event, we recorded the presence or absence of at least one copepod harboring an infective (L3) larva within the sampled volume. A copepod was considered detectable if it was spatially located inside the sampling sphere and contained at least one L3 larva at that time point. The primary modeling outcome was the detection probability, calculated as the proportion of samples in which at least one infected copepod was identified, out of 1,000 simulated sampling iterations. This detection probability served as a quantitative measure of surveillance sensitivity, reflecting the likelihood of detecting infected copepods via routine water sampling across varying ecological contexts and sampling intensities relevant to environmental surveillance in endemic settings.

### Baseline model

The baseline simulation was implemented using fixed parameter values for five key ecological and environmental variables (Table 1): larval diffusion coefficient (*D*_*c*_), larval lifespan (*T*_*L*_), copepod density (*ρ*), edge preference (γ_edge_), ingestion probability (*P*_*ingest*_), and pond radius (*R*). These parameters were selected for their biological relevance to *D. medinensis* transmission and environmental detection dynamics. All other model conditions—including pond geometry, copepod and larval behavior, and water sampling—were defined as described above.

**Table 1.** Parameters in sensitivity analysis. Baseline values are underlined.

| Parameter Name | Notation | Level |  |  |  |  | Units |
| --- | --- | --- | --- | --- | --- | --- | --- |
|  |  | 1 | 2 | 3 | 4 | 5 |  |
| **Larval diffusion coefficient** | $D_c$ | <u>0.0035</u> | 0.007 | 0.010 | 0.015 | 0.020 | m<sup>2</sup> / day |
| **Average Larval lifespan** | $T_L$ | 1 | 3 | <u>5</u> | 7 | 9 | Days |
| **Copepod density** | $\rho$ | 1 | 3 | <u>10</u> | 15 | 20 | Individual /<br>L |
| **Edge preference** | $\gamma_{\text{edge}}$ | 0.1 | 0.3 | <u>0.5</u> | 0.7 | 0.9 | Unitless |
| **Ingestion probability** | $P_{\text{ingest}}$ | 0.01 | 0.05 | <u>0.1</u> | 0.3 | 0.5 | Per<br>encounter |
| **Radius of pond** | $R$ | 1 | 2 | 3 | 4 | <u>5</u> | Meters |

### Sensitivity analysis

#### One-at-a-time (OAT)

To evaluate the impact of ecological and environmental uncertainty on surveillance outcomes, we conducted a one-at-a-time (OAT) sensitivity analysis by systematically varying each of six key parameters—*D*_*c*_, *T*_*L*_, *ρ*, γ_edge_, *P*_*ingest*_, and *R*— across five biologically plausible levels while holding the others at their baseline values (Table 1). Detection probability time series were analyzed to quantify the independent effect of each parameter, identifying which factors most strongly influenced environmental surveillance sensitivity.

### Interaction Effects

To evaluate how combinations of parameters jointly affect surveillance outcomes, we performed a pairwise interaction analysis using a full-factorial design. Simulations were conducted for all unique combinations of the six key parameters (Table 1), with model output defined as the cumulative number of days on which at least one L3-infected copepod was detected across 1,000 randomly distributed 24-liter water samples per day. A full-factorial ANOVA was then applied to these outputs to identify significant second order (pairwise) interaction effects. The strength and structure of these interactions were visualized using a chord diagram, where thicker chords represent stronger interactions between parameters. This analysis allowed identification of parameter combinations that most strongly influenced environmental surveillance performance under complex ecological conditions.

### Multi-way

Building on previous sensitivity analyses, we identified larval diffusion coefficient (*D*_*c*_), copepod density (*ρ*), and pond radius (*R*) as key drivers of surveillance sensitivity. To assess their combined effects, we performed a grid-based multi-way (factorial) sensitivity analysis, systematically varying *D*_*c*_, *ρ*, and *R* across their respective ranges (Table 1) and simulating all possible parameter combinations. For each scenario, model output was defined as the cumulative probability of detecting at least one L3-infected copepod across repeated water sampling events. This approach allowed us to comprehensively evaluate how interactions among these core ecological and environmental parameters shape detection probability, providing deeper insight into the mechanisms governing environmental surveillance outcomes in complex pond systems.

### Sampling strategy implementation

To evaluate how sampling design influences surveillance sensitivity, we examined two complimentary aspects of sampling strategy. First, we systematically varied the number and size of water samples, while maintaining equivalent total sampling volumes across different combinations, allowing comparison of detection probabilities between strategies using many small samples versus fewer large samples. These analyses were conducted using baseline values for all other parameters and repeated across a range of larval diffusion coefficients (*D*_*c*_) to assess interactions between sampling configuration and larval dispersal dynamics.

Second, we explored the effects of spatial targeting by varying the range over which sampling locations were distributed relative to the modeled larval release point. Although the exact location of larval release would be unknown in practice, examining detection probability as a function of distance from the release point is informative for understanding how spatially uneven larval distributions can create heterogeneity in detection. Sampling locations were randomly generated within specified distance ranges and constrained within pond boundaries, allowing quantification of how detection probability declines with increasing distance from areas of larval introduction.

Third, to address surveillance performance under realistic spatial uncertainty, we evaluated sampling strategies that do not rely on knowledge of the larval release location. In this analysis, sampling locations were defined relative to the pond shoreline, using fixed-width sampling bands extending inward from the pond edge. These shoreline-based sampling strategies were evaluated across varying levels of copepod edge preference, which governs the degree to which copepods aggregate near pond margins versus being evenly distributed throughout the water body. By combining shoreline-targeted sampling with different copepod movement behaviors and larval diffusion rates, this analysis assessed how detection probability depends on the alignment between sampling design and predictable ecological structure, rather than on knowledge of localized larval hotspots. Together, these analyses were designed to identify operationally relevant sampling approaches that may improve environmental surveillance for Guinea worm in heterogeneous aquatic environments.

## Results

### Baseline model

The baseline simulation produced distinct temporal and spatial patterns in copepod infection dynamics (Fig 3A). Copepods with early-stage (L1/L2) larvae peaked shortly after larval release and declined thereafter, while copepods harboring infective stage (L3) larvae emerged later, with peak abundance occurring around day 23. Consequently, the probability of detecting at least one L3-infected copepod in a water sample increased sharply during days 20–45, reaching a maximum near day 30 before dropping as infected hosts became rare (Fig 3B).

**Fig 3.**
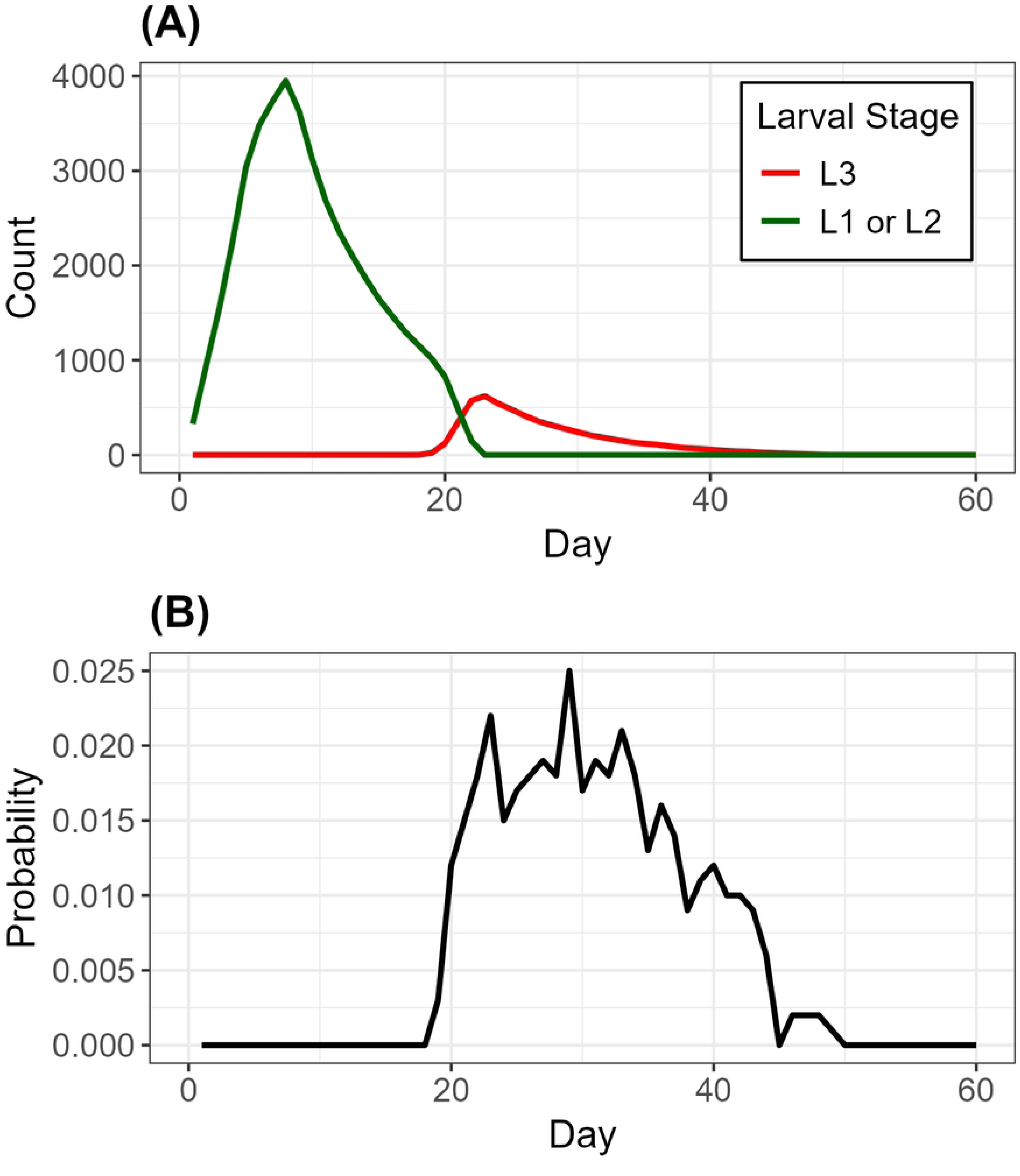
Temporal infection and detection dynamics. **(A)** Temporal dynamics of copepod infection. The number of copepods carrying early-stage (L1/L2) larvae (green) peaks within the first two weeks post-release, then declines as larvae mature and hosts die. The number of copepods with infective-stage (L3) larvae (red) peaks around day 30. **(B)** Daily probability of detecting at least one copepod with L3s in a 24-liter water sample.

Spatially infected copepods were heterogeneously distributed. A representative snapshot (Fig 2) shows that L3-infected individuals were most concentrated at intermediate distances from the larval release point, reflecting the combined effects of larval dispersal, infection-induced mortality near the release site, and copepod movement toward the pond periphery. These baseline patterns suggest that environmental surveillance sensitivity is temporally constrained and strongly influenced by the spatial ecology of infected copepods within the pond environment. Accordingly, these baseline parameters served as a reference for interpreting subsequent analyses of sensitivity and sampling-strategies.

### Sensitivity analysis

#### One-at-a-time (OAT)

Figure 4 illustrates the temporal effects of each parameter on the probability of detecting at least one L3-infected copepod in a 24-liter water sample. The detection probability was highly sensitive to larval diffusion coefficient (*D*_*c*_), larval lifespan (*T*_*L*_), and copepod density (*ρ*). The effect of larval diffusion was non-linear: detection was maximized at intermediate larval diffusion coefficient values, with both low and high diffusion reducing contact between larvae and hosts. Longer larval lifespans shifted detection curves later and increased their amplitude and duration, as extended larval availability allowed sustained ingestion over time. Increasing copepod density raised detection probability initially by increasing encounter rates; however, gains diminished at higher densities, reflecting saturation of the binary detection metric once the probability of encountering at least one infected copepod approached its upper bound. Edge preference (γ_edge_) affected detection by modulating spatial overlap. Stronger edge bias reduced overlap with centrally located samples, suppressing detection probability and underscoring the dependence of surveillance sensitivity on aligning sampling location with biologically relevant copepod distributions. Ingestion probability (*P*_*ingest*_) had limited impact except at unrealistically low values, indicating that ingestion efficiency is not a primary limiting factor under typical ecological conditions. Lastly, pond radius (*R*) exhibited a strong negative effect—larger ponds diluted agent densities, decreasing encounter frequency and reducing both peak magnitude and detection window.

**Fig 4.**
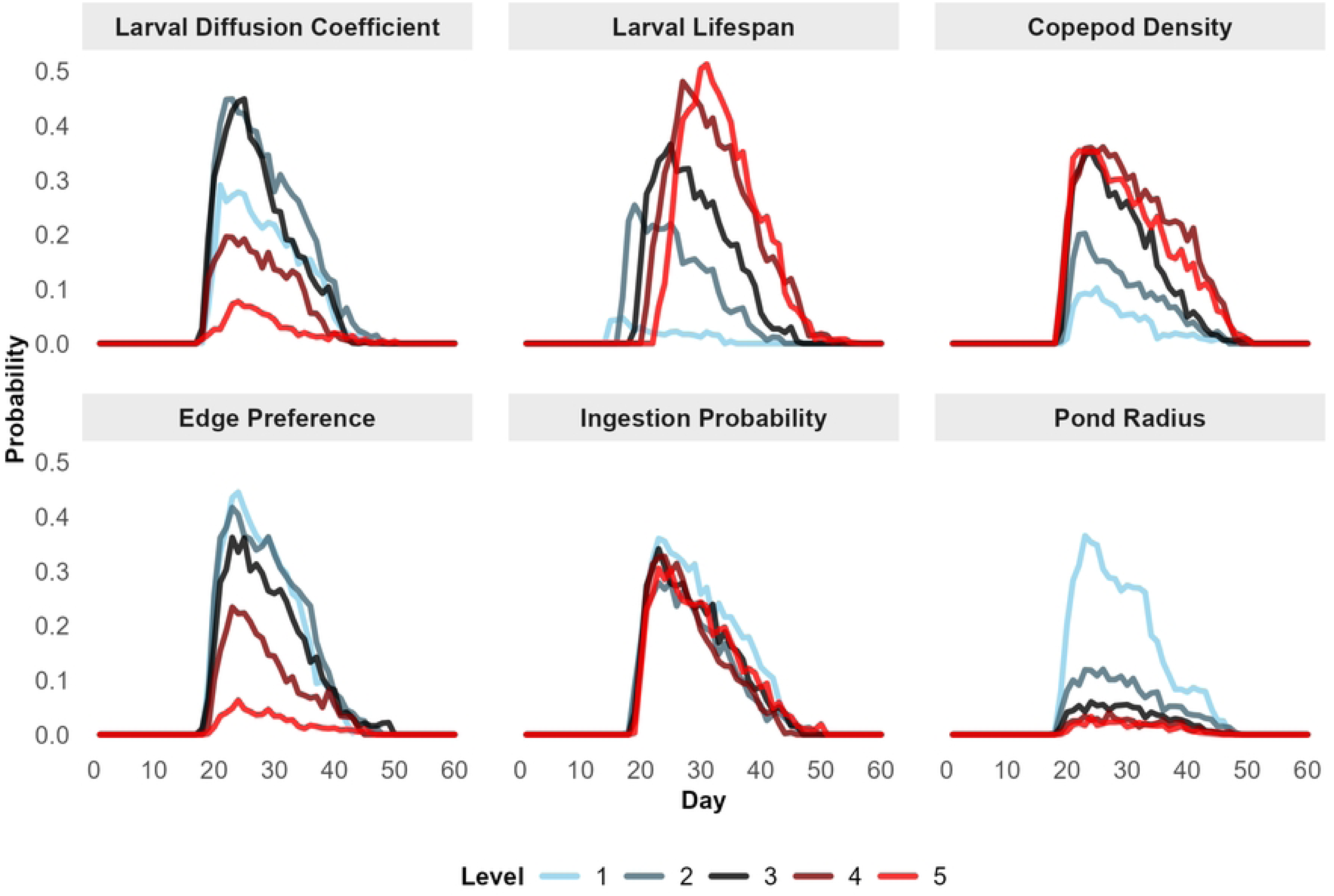
Temporal effects of individual parameters on detection probability in one-at-a-time (OAT) sensitivity analysis. Each panel shows detection probability over time for one parameter varied across five levels, which is shown in Table 1, with others held constant.

### Interaction Effects

Analysis of pairwise interactions revealed that model outcomes were shaped not only by individual parameter effects but also by strong interdependencies between key ecological drivers. As visualized in the chord diagram (Fig 5), the larval lifespan (*T_L_*) exhibited prominent interactions with multiple parameters, including copepod density (*ρ*), larval diffusion coefficient (*D_c_*), and pond radius (*R*). These interactions suggested that the benefit of extended larval survival was contingent on sufficient dispersal and encounter opportunities, which are in turn influenced by host density and spatial extent. The interaction between *D_c_* and *R* indicated that habitat spatial scale modulates how larval diffusion influenced detection probability. Similarly, the strong *T_L_***–** ρ interaction indicates that longer-lasting larvae contribute to detection sensitivity primarily when host density was adequate to support ingestion and development.

**Fig 5.**
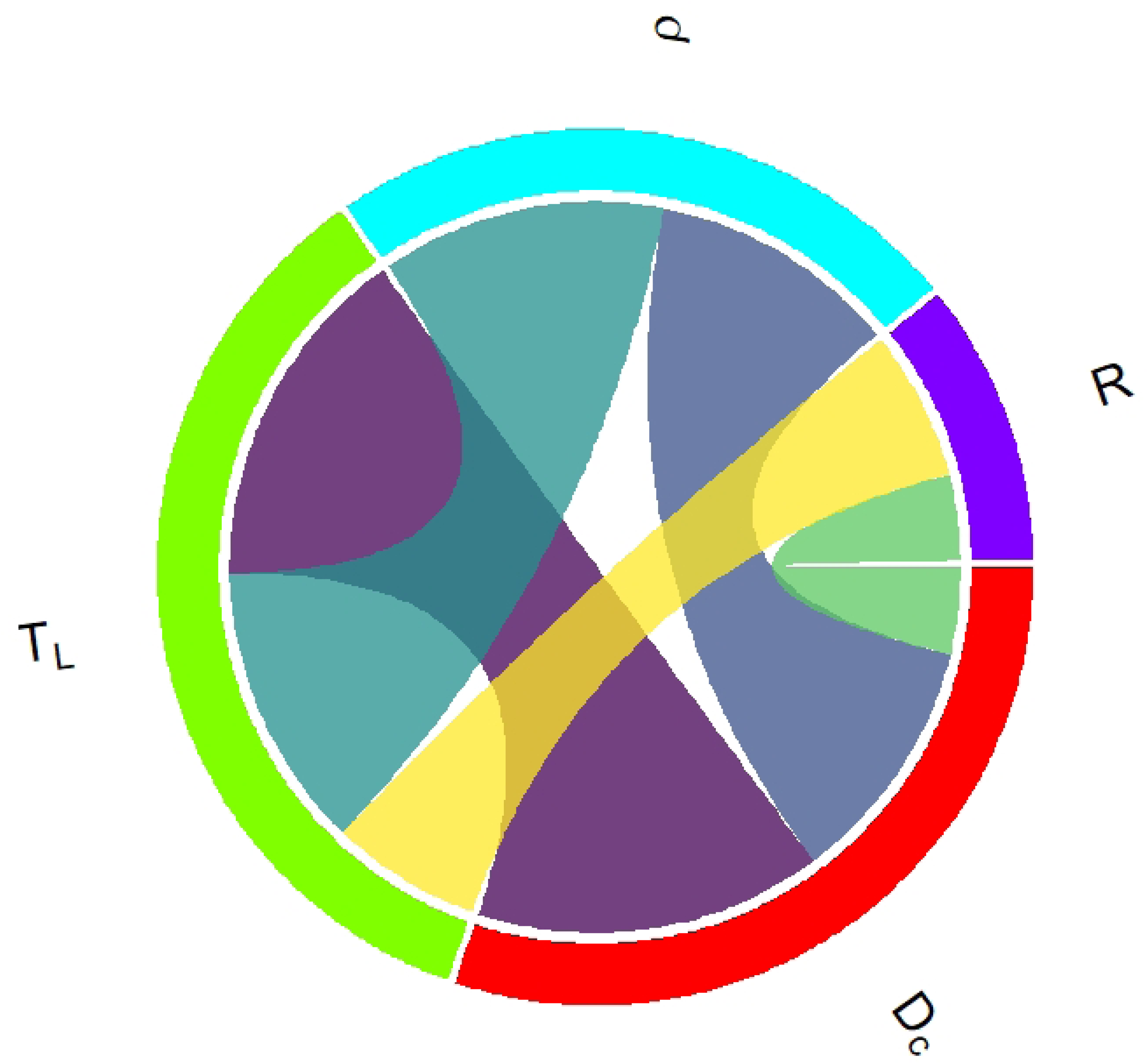
Chord diagram depicting the strongest pairwise parameter interactions affecting detection probability of L3-infected copepods (top 25% by variance explained from the ANOVA). Each arc corresponds to a model parameter—larval diffusion coefficient (*D*_*c*_), larval lifespan (*T*_*L*_), copepod density (*ρ*), and pond radius (*R*)—and the width of each connecting ribbon reflects the magnitude of the interaction. Symbols correspond to parameter notations, highlighting interacting parameter pairs that jointly influence model behavior.

### Multi-way

Based on the one-at-a-time and pairwise interaction analyses, larval diffusion coefficient (*D*_*c*_), copepod density (*ρ*), and pond radius (*R*) were identified as the dominant drivers of detection probability; accordingly, the multi-way analysis focused on interactions among these parameters. Detection probability exhibited distinct patterns across combinations of *D*_*c*_, *ρ*, and *R* (Fig 6). As in the baseline model, copepod abundance scaled with pond volume while larval numbers were fixed, so the larval-to-copepod ratio declined with increasing pond size. Under low copepod density (*ρ* = 1), detection probability increased slightly with *D*_*c*_ in medium (*R* = 3 *m*) and large (*R* = 5 *m*) ponds, likely because greater dispersal helped the fixed larval pool reach a wider portion of the copepod population in larger habitats. In small ponds (*R* = 1 *m*), detection peaked at the lowest *D*_*c*_before declining, indicating that limited dispersal concentrated larvae near sampling zones, whereas higher *D*_*c*_dispersed them more evenly and reduced local larval densities and detection probability.

**Fig 6.**
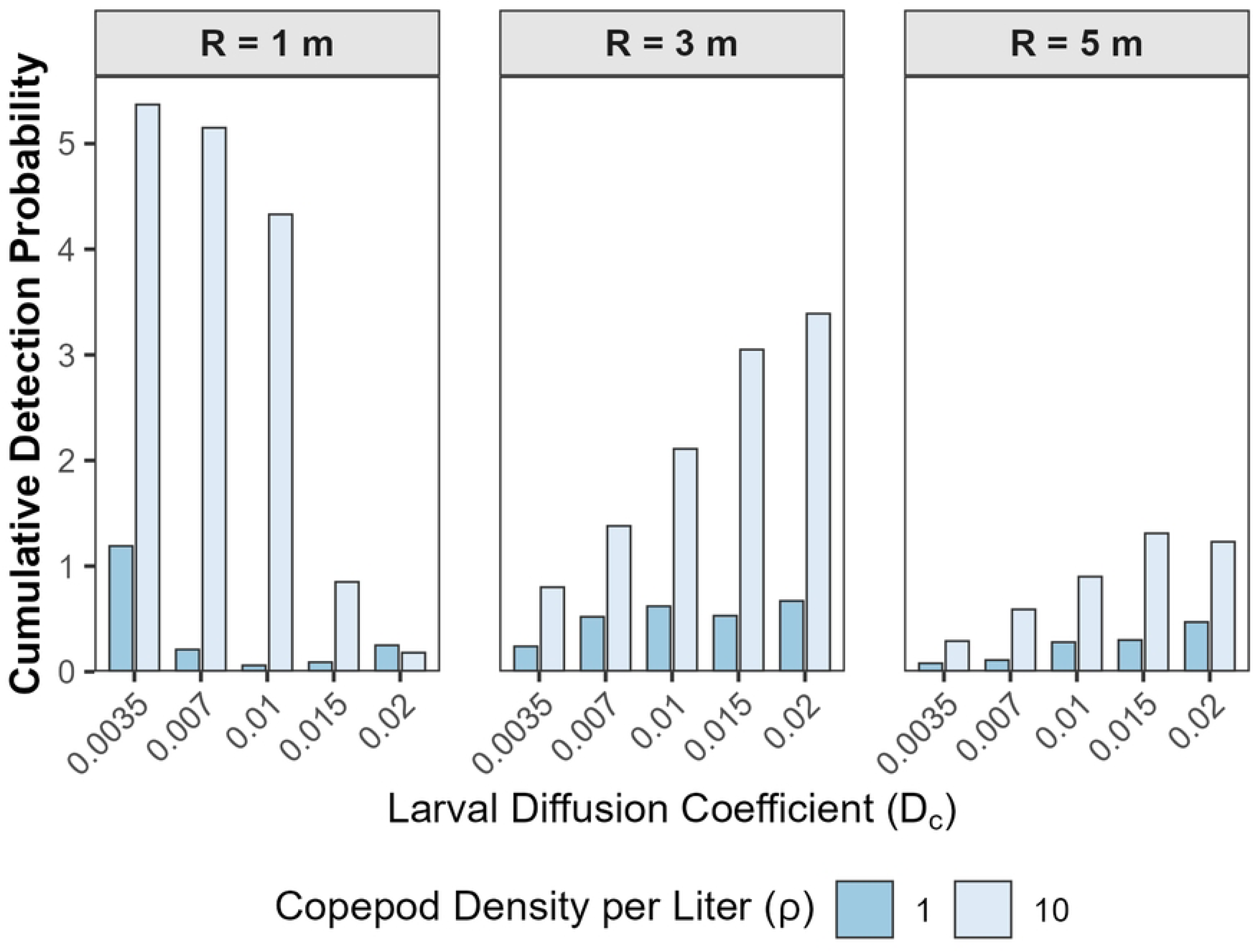
Effects of pond radius, larval diffusion coefficient, and copepod density on cumulative detection probability. The y-axis shows the sum of daily detection probabilities over the 60-day simulation period, representing the cumulative probability of detecting at least one L3-infected copepod in water samples. Each panel corresponds to a different pond radius (R), with bars indicating cumulative detection probability for two copepod densities (ρ) (1 and 10 per liter) across a range of larval diffusion coefficients (*D*_*c*_). This metric quantifies the overall likelihood of surveillance success under varying ecological scenarios.

At high copepod density (*ρ* = 10), detection probability in small ponds decreased monotonically with increasing *D*_*c*_. In this setting, high copepod abundance led to rapid ingestion of larvae near the sampling zone at low *D*_*c*_, while increased dispersal spread larvae more broadly, lowering their local density despite abundant copepods. In contrast, in medium and large ponds, the larger total number of copepods meant that larvae were initially concentrated near the release point, but many copepod hosts were located farther away. Higher *D*_*c*_allowed larvae to disperse across the pond and reach a greater fraction of the host population, which increased the probability of detecting infected copepods. Overall, these results indicated that the interaction between larval movement, host density, and spatial scale was strongly shaped by the fixed larval input relative to pond volume. Although larval dispersal mechanisms themselves were independent of pond size, their effects on detection differed by spatial scale: in small ponds, dispersal could reduce local larval concentrations near sampling zones, whereas in larger ponds it facilitated encounters with a more spatially distributed copepod population.

### Sampling strategies

The relationship between sampling configuration and detection probability depended on the effective larval diffusion rate, which in this model represents differences in environmental mixing rather than changes in larval behavior (Fig 7). At low and intermediate diffusion coefficients (*Dc*=0.0035 and 0.01), cumulative detection probability increased both with greater total sampling volume and with a larger number of samples, even when total volume was held constant. This suggested that, under these diffusion regimes, distributing sampling effort across more locations improved the chances of detecting infected copepods. In contrast, at the highest diffusion coefficient (*Dc*=0.02), detection probability remained relatively uniformly low and showed little sensitivity to either total sampling volume or number of samples. These results indicate that spatially distributed sampling may improve environmental surveillance performance when larvae remain spatially concentrated within ponds.

**Fig 7.**
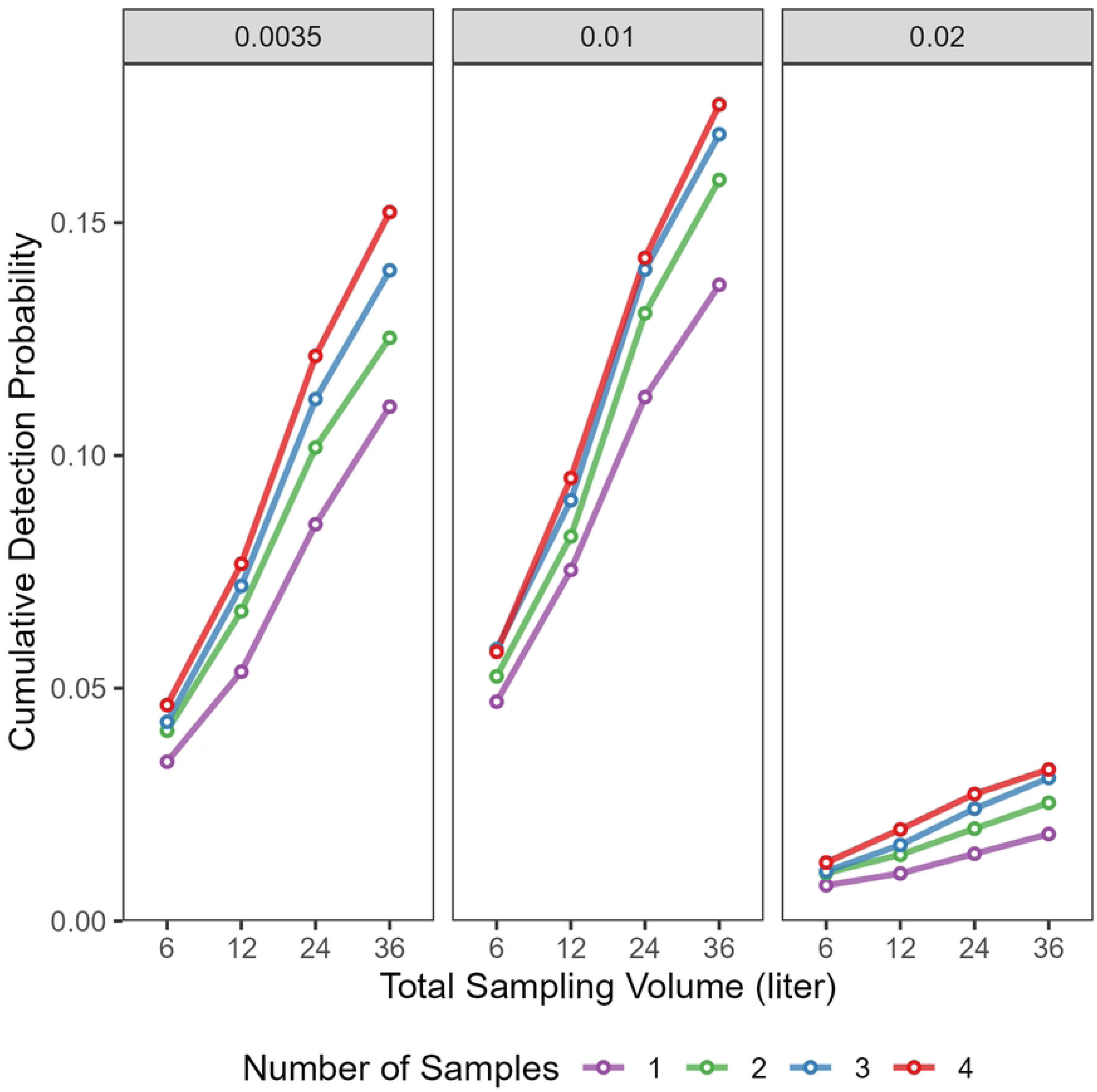
Effects of sampling configuration and larval diffusion on cumulative detection probability. Cumulative probability of detecting at least one L3-infected copepod over the 60-day simulation period as a function of total sampling volume (x-axis) and number of samples (color lines), shown for three larval diffusion coefficients (Dc=0.0035, 0.01, and 0.02). Each line corresponds to a different number of samples (1–4), with volume per sample accommodated to keep total sampling volume held constant across combinations.

Detection probability was jointly influenced by sampling distance from the larval release point and the effective larval diffusion coefficient (Fig 8). At short sampling distances from the pond edge (0.4 and 0.7 m), detection events were rare and primarily occurred only at the lowest larval diffusion coefficient (*D_c_*= 0.0035 m²/d), corresponding to scenarios in which larvae remain concentrated near entry points. As sampling distance increased, detection probability for this lowest diffusion coefficient rose sharply, peaking at approximately 1 m, before declining at greater distances as larvae remained localized near the source. In contrast, scenarios with higher diffusion coefficients (*D_c_* = 0.01, 0.015, and 0.02 m²/d) exhibited low detection probabilities near the release point but increased detection at sampling distances of 2—3 m from the pond edge, consistent with broader larval redistribution across the waterbody. These results suggest that the spatial scale at which sampling was most effective depends on extent of larval dispersal within the aquatic environment.

**Fig 8.**
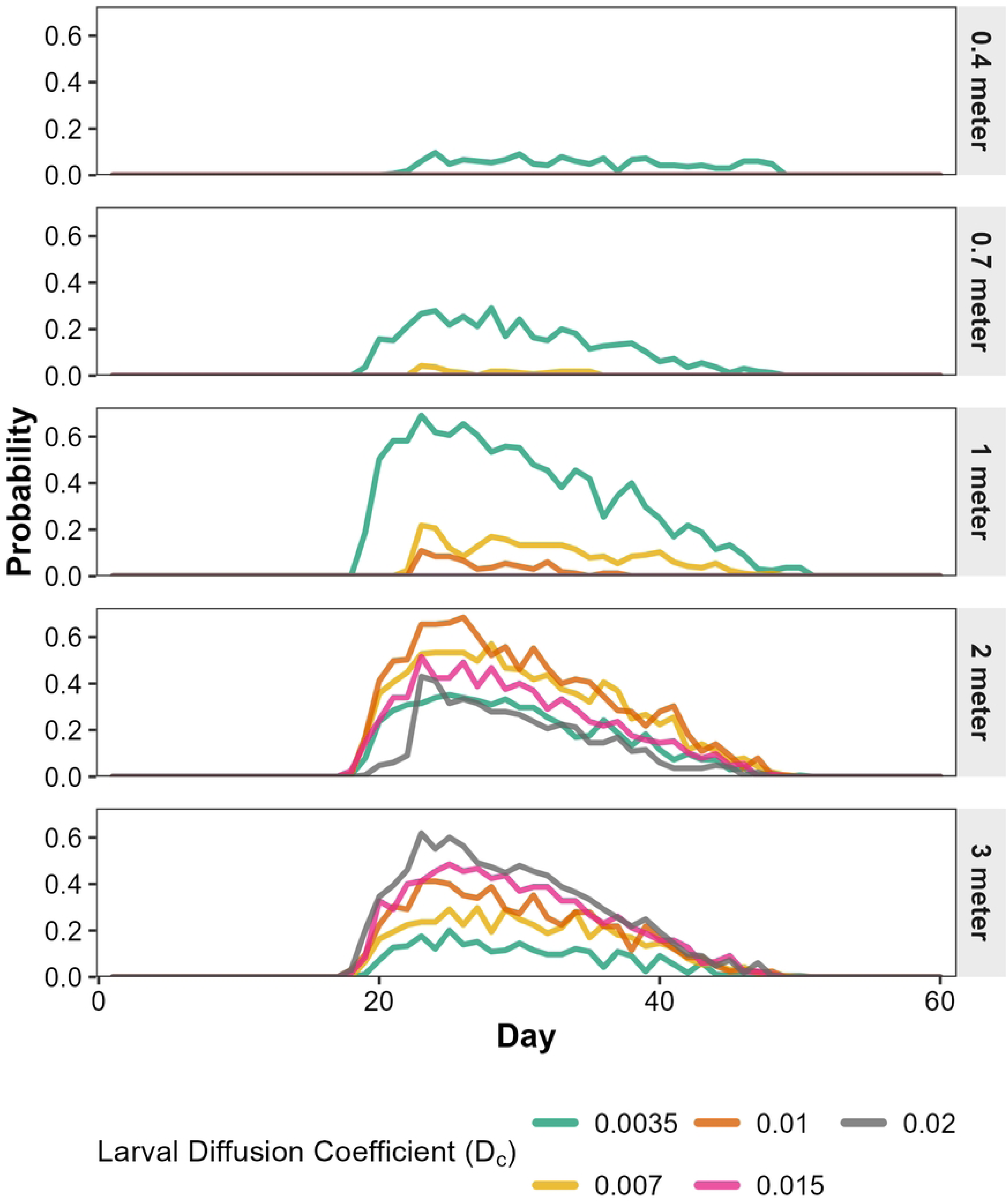
Influence of sampling distance and larval diffusion on temporal detection probability. Probability of detecting at least one L3-infected copepod is shown over time for varying radial distances from the larval release point (panels, 0.4–3 meters) and larval diffusion coefficients (Dc=0.0035 to 0.02; colored lines). For each scenario, sampling locations were randomly distributed within the defined distance from the release site at the pond edge.

Detection probability varied strongly with the interaction between copepod edge preference, shoreline-based sampling band width, and larval diffusion (Fig 9). When copepods were weakly edge-biased, detection probability was relatively insensitive to sampling band width, with comparable detection trajectories observed across narrow and broad shoreline bands. In contrast, as copepod edge preference increased, detection probability became increasingly concentrated in narrow shoreline sampling bands, with substantially higher detection achieved when sampling was restricted to locations close to the pond boundary. Expanding the sampling band inward resulted in progressive declines in detection probability under strong edge preference, reflecting reduced overlap between sampling locations and copepod aggregation. Across all scenarios, higher larval diffusion coefficients increased overall detection probability, but this effect was most pronounced when shoreline-based sampling aligned with copepod spatial distribution.

**Fig 9.**
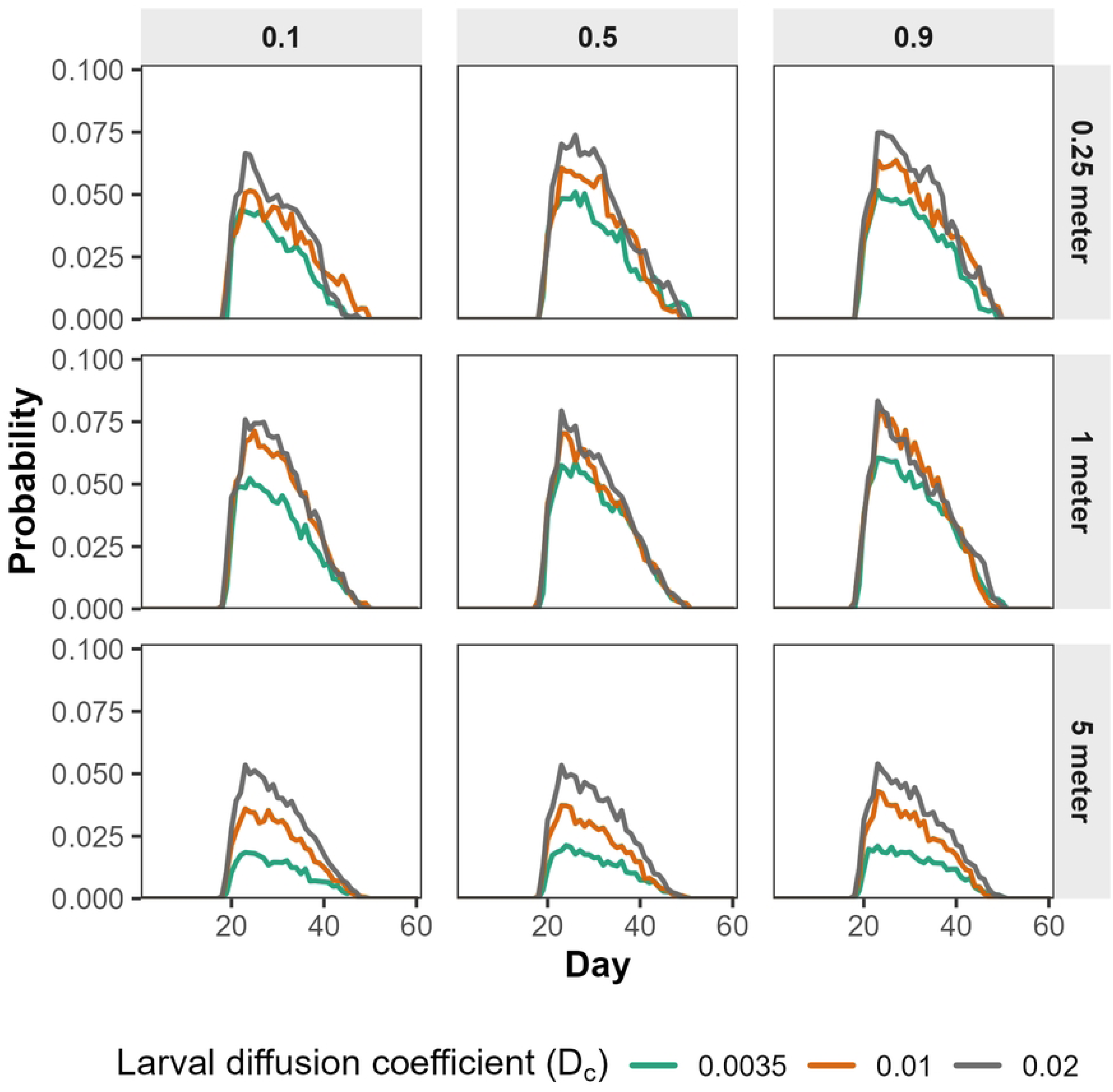
Detection probability under shoreline-based sampling across three levels of copepod edge preference and sampling range. Columns represent copepod edge preference (*γ* = 0.1, 0.5, 0.9), indicating increasing tendency of copepods to aggregate near the pond edge. Rows represent shoreline-based sampling bands extending inward from the pond edge (0.25 m, 1 m, and 5 m, where 5 m corresponds to sampling across the entire pond). Colored lines indicate larval diffusion coefficients (*D*_*c*_).

Temporal patterns were characterized by an early rise and subsequent decline in detection probability, with peak magnitude and persistence dependent on both copepod behavior and sampling design. Together, these results demonstrated that surveillance performance under spatial uncertainty depends strongly on alignment between ecological structure and shoreline-based sampling design.

## Discussion

This study used a 3D, spatially explicit, ABM to examine how ecological processes shape the probability of detecting *D. medinensis*–infected copepods through environmental water sampling. Rather than identifying a single dominant driver, the model demonstrated that surveillance sensitivity emerges from the interaction among larval diffusion, copepod density, pond size, and sampling strategy. These interactions generated nonlinear and context-dependent detection patterns, highlighting that surveillance outcomes cannot be inferred from any single ecological parameter in isolation. By explicitly representing host–parasite interactions in a mechanistic framework (23), this model provides a framework for interpreting environmental surveillance results and translating ecological complexity into programmatically meaningful guidance for Guinea worm surveillance.

A central finding of this work is that detection of L3-infected copepods is intrinsically constrained to a narrow spatiotemporal window. Detection probability peaked after ingested larvae matured to the infective stage but declined rapidly thereafter due to larval mortality, infection-induced copepod death, and continued dispersal. Even under favorable ecological conditions, this window was brief relative to the total simulation period, indicating that environmental surveillance has inherently limited sensitivity and is most likely to succeed during a short interval when larval abundance and host encounter rates are simultaneously elevated (24). These findings suggest that the timing of water sampling may strongly influence surveillance performance in endemic settings. This temporal constraint has direct implications for Guinea worm surveillance programs. Negative water samples do not provide definitive evidence that larvae are absent. When the timing of larval introduction into waterbodies is uncertain—as is typical in field settings—effective surveillance requires repeated sampling over time to capture transient windows of detectability (25,26).

Detection was further constrained by spatial heterogeneity in infected copepod distribution. Infected copepods formed clusters shaped by larval dispersal and copepod movement behavior, with early infections concentrated near larval entry points and later infections occurring at greater distances as larvae dispersed and infected copepods died (18). Similar spatial and temporal heterogeneity has been observed in other aquatic disease systems, including schistosomiasis and malaria vector ecology. This underscores the need for surveillance strategies that explicitly account for clustering rather than relying on uniform sampling assumptions (29,30). These results suggest that spatially uniform water sampling may overlook localized areas of elevated transmission risk. Failure to account for these spatial and temporal constraints may lead to false confidence in elimination status, particularly during the final stages of eradication.

Larval diffusion emerged as a key driver of detection probability, but its effects were strongly nonlinear. Detection was maximized at intermediate diffusion rates that allowed larvae to disperse beyond initial entry points while maintaining sufficient local concentrations to support ingestion by copepods. At low diffusion, larvae remained spatially constrained, limiting encounters with hosts, whereas at high diffusion larvae became diluted across the waterbody, reducing encounter rates despite broader distribution. These findings indicate that environmental mixing may influence surveillance sensitivity in contrasting ways depending on the extent of larval dispersal within a pond. Surveillance is therefore most effective in moderately mixed ponds that redistribute larvae without fully homogenizing them (19).

The nonlinear effects of larval diffusion on detection probability were further modulated by pond size through changes in the larval-to-copepod ratio. Because larval input was fixed while copepod abundance scaled with pond volume, larger ponds experienced lower local larval densities and reduced encounter rates. Detection probability declined accordingly, particularly at low copepod densities. Importantly, the influence of larval diffusion depended on spatial scale: in small ponds, increased diffusion often reduced detection by dispersing larvae away from sampling zones, whereas in larger ponds diffusion improved detection by enabling larvae to reach a more spatially distributed copepod population. Copepod density partially mitigated these effects, emphasizing that surveillance outcomes are governed by interactions with ecological drivers rather than isolated factors (27,28).

Spatial targeting of sampling effort can substantially influenced detection sensitivity under uncertainty. Across many scenarios, detection probability was highest when sampling locations aligned with predictable copepod aggregation near pond margins. Shoreline-based sampling strategies consistently outperformed uniform or whole-pond sampling when copepods exhibited even moderate edge preference (15), suggesting that prioritizing sampling near pond edges and sites of frequent human or animal water contact may improve surveillance efficiency when larval entry points are unknown. Sampling configuration also influenced surveillance outcomes. When total sampling volume was held constant, distributing effort across multiple spatial locations modestly but consistently improved detection probability in smaller or weakly mixed ponds where infected copepods remained spatially clustered. This finding supports surveillance strategies that prioritize spatial replication over maximizing single-sample volume, particularly when laboratory capacity or field resources are constrained. In contrast, under highly mixed conditions or in larger ponds, the marginal benefit of additional spatial replication declined as infected copepods became more broadly dispersed throughout the waterbody (31). Collectively, these findings indicate that environmental surveillance performance depends strongly on how well sampling strategies align with underlying ecological structure and spatial heterogeneity within aquatic habitats.

Observable pond characteristics—including size, apparent mixing, and copepod abundance—can inform both surveillance design and interpretation. Larger ponds and highly mixed systems are likely to yield low detection probabilities even when transmission persists, whereas moderately mixed, smaller waterbodies provide more favorable conditions for environmental detection. Incorporating such contextual information into surveillance planning may improve targeting efficiency and reduce overinterpretation of negative water samples, without requiring precise knowledge of larval release timing or location (32).

More broadly, this modeling framework demonstrates how mechanistic, spatially explicit models can support disease elimination programs by translating ecological complexity into actionable surveillance guidance (33). Although the model simplifies natural systems by excluding hydrological flow, seasonal dynamics, interspecific copepod variation, and animal reservoirs, it captures key constraints that shape environmental detection. Future work incorporating these processes and field validation data will further refine surveillance expectations, but the present results already highlight that effective Guinea worm environmental surveillance requires adaptive, context-aware strategies rather than uniform sampling designs.

### Conclusion

Ongoing development of water-based assays capable of detecting *D. medinensis* analytes offers new opportunities for environmental surveillance, but their effective deployment requires an understanding of field-level detection sensitivity. This study demonstrated the utility of 3D ABM for examining how ecological processes shape detection of *D. medinensis* in aquatic environments. By showing how the interplay of larval diffusion, pond size, copepod density, and sampling design constrains detection probability, the model provides practical guidance for refining sampling strategies for Guinea worm surveillance and for interpreting environmental surveillance outcomes. Future work incorporating hydrological processes, variable copepod population dynamics, interspecific behavioral variation, and potential definitive animal hosts will further improve ecological realism. As eradication efforts enter their final phase, integrating mechanistic modeling into surveillance planning can help improve targeting of limited surveillance resources and support efforts to identify residual transmission.

## Acknowledgments

We thank the Guinea Worm Ecology Working Group for valuable discussions and feedback. We are especially grateful to Zachary Wagner and Jeannette Yen for their guidance on parameterization. We also thank the reviewers from The Carter Center, particularly Kathryn Schaber and Jessica van Loben Sels, for their constructive comments and insights, which helped improve the clarity and relevance of this work.

## Supporting information

**S1 File. Operational guidance for water sampling to detect Dracunculus medinensis–infected copepods.** Model-informed recommendations on sampling design, spatial targeting, and temporal repetition to improve surveillance under ecological uncertainty.

